# Concurrent Guidance of Attention by Multiple Working Memory Items: Behavioral and Computational Evidence

**DOI:** 10.1101/629378

**Authors:** Cherie Zhou, Monicque M. Lorist, Sebastiaan Mathôt

**Affiliations:** Department of Experimental Psychology, University of Groningen; Faculty of Medical Sciences, Biomedical Sciences of Cells & Systems, Neurosciences, Neuroimaging Center, University of Groningen, The Netherlands

**Keywords:** visual working memory, visual search, attentional guidance, attentional capture, drift diffusion

## Abstract

During visual search, task-relevant representations in visual working memory (VWM), known as attentional templates, are assumed to guide attention. A current debate concerns whether only one (Single-Item-Template hypothesis, or SIT) or multiple (Multiple-Item-Template hypothesis, or MIT) items can serve as attentional templates simultaneously. The current study was designed to test these two hypotheses. Participants memorized two colors, prior to a visual-search task in which the target and the distractor could match or not match the colors held in VWM. Robust attentional guidance was observed when one of the memory colors was presented as the target (reduced response times [RTs] on target-match trials) or the distractor (increased RTs on distractor-match trials). We constructed two drift-diffusion models that implemented the MIT and SIT hypotheses, which are similar in their predictions about overall RTs, but differ in their predictions about RTs on individual trials. Critically, simulated RT distributions and error rates revealed a better match of the MIT hypothesis to the observed data than the SIT hypothesis. Taken together, our findings provide behavioral and computational evidence for the concurrent guidance of attention by multiple items in VWM.

**Significance statement:** Theories differ in how many items within visual working memory can guide attention at the same time. This question is difficult to address, because multiple- and single-item-template theories make very similar predictions about average response times. Here we use drift-diffusion modeling in addition to behavioral data, to model response times at an individual level. Crucially, we find that our model of the multiple-item-template theory predicts human behavior much better than our model of the single-item-template theory; that is, modeling of behavioral data provides compelling evidence for multiple attentional templates that are simultaneously active.

## Simultaneous Guidance by Multiple Attentional Template

Internal representations of task-relevant information, or *attentional templates*, stored in visual working memory (VWM), guide attention in visual search (Bundesen, 1990; Bundesen et al., 2005). For example, when you are looking for a chocolate cake, all dark items in a bakery will be more likely to draw your attention. The biased-competition framework (Desimone, 1998) states that VWM leads to pre-activation of memorized features in visual cortex. In this example, when you keep the color of a chocolate cake in VWM, neurons in color-selective areas that represent this color become pre-activated. And later, when the color is actually perceived, this pre-activation leads to an enhanced neural response, which at the behavioral level results in attention being drawn towards chocolate-cake-like objects. In other words, VWM contents guide attention towards memory-matching items in a top-down manner to optimize visual search (Chelazzi, Miller, Duncan, & Desimone, 1993; Chelazzi, Duncan, Miller, & Desimone, 1998).

Although multiple representations can be maintained in VWM simultaneously, there is ongoing debate about the number of VWM items that can simultaneously serve as attentional templates. The Single-Item-Template hypothesis (SIT; Houtkamp & Roelfsema, 2006; Olivers, Peters, Houtkamp, & Roelfsema, 2011) proposes a functional division within VWM: While one item actively interacts with visual processing to guide attentional selection towards matching items, other items are shielded from visual sensory input, and thus cannot guide attention.

Studies demonstrating a switch cost between templates are often interpreted as evidence for the SIT model. In a study by Dombrowe, Donk, and Olivers (2011), participants made a sequence of two eye movements towards two spatially separated target items which were indicated by arrows. In the switch condition, the two targets had different colors, and thus required a switch between two templates; in the no-switch condition, both targets had the same color, and thus required only one attentional template. Crucially, eye movements that were correctly aimed at the second target were delayed by about 250ms – 300ms in the switch condition, compared to the no-switch condition. This cost associated with switching between templates is in line with the SIT hypothesis, suggesting that only one template can be active at one time.

In contrast to the SIT hypothesis, the Multiple-Item-Template (MIT) hypothesis suggests that multiple VWM items can guide attention simultaneously (Beck et al., 2012), although holding multiple items in VWM would reduce the memory quality of each item, thus reducing memory-driven guidance (Bays & Husain, 2008; Kristjánsson & Kristjánsson, 2018). As Kristjánsson et al. (2018) point out, even if multiple VWM items *can* guide attention simultaneously, this does not mean that they always do; specifically, they propose that multiple VWM items guide attention at the same time only when this is needed for the task. The MIT hypothesis builds on research suggesting that there is no unitary spotlight of attention, but rather that attention can be divided (Eimer & Grubert, 2014), in this case, across multiple memory-matching items.

Recent work by (Beck & Hollingworth, 2017) supported the MIT hypothesis. In their experiment (a saccadic sequential search task), participants first saw a cue that consisted of two colors (e.g., red and blue), followed by two pairs of colored objects, presented one pair at a time. The first pair always contained one non-matching distractor (e.g., yellow) and one object that matched one of the cued colors (e.g., red); participants fixated this cue-matching object. In the second pair, the cue-matching object from the first pair was presented either with a new non-matching distractor (e.g., green), or with an object that matched the remaining cued color (blue). In the latter case, participants were free to select either object. Critically, when participants were free to select either the first- or the second-cued color in the second pair, the selection probability of the first cued color was substantially reduced: They were about as likely to first select red and then blue, as they were to select red twice. In other words, even though participants presumably had an active search template for the first-cued color, the second-cued color was able to compete with it. This competition between the two cue-matching objects suggests that both templates were maintained in an active state in VWM.

However, when looking at behavioral evidence comparing the SIT and MIT hypothesis (e.g., (Hollingworth & Beck, 2016; van Moorselaar et al., 2014), it is difficult to distinguish between the two hypotheses by only observing average RTs across trials. A more powerful way to distinguish the underlying cognitive processes is by analyzing RT distributions, an approach that has been used successfully in previous studies. For example, (Chetverikov A et al., 2017; Chetverikov et al., 2016)looked at RT distributions to test how different properties of previously observed distractor distributions (e.g., shape) influence search times. And (Sung, 2008) analyzed RT distributions for displays of different set sizes to distinguish parallel from serial mechanisms in visual selection. Following this approach, in the current study, we compared not only the average RTs, but also the RT distributions of trials in different conditions under the SIT and the MIT hypothesis. Critically, we simulated individual trials based on the predictions of two hypotheses by means of a drift-diffusion model (Ratcliff & McKoon, 2008) and compare the simulated data to the obtained data. We implemented a visual-search task based on the additional-singleton paradigm (Theeuwes, 1992). Participants first kept two colors in working memory, after which they searched for a colored target shape among a colored distractor shape and, in one experiment (Experiment 1), a grey distractor shape. The color of the target and the (colored) distractor was manipulated to match or not match the memorized colors.

Overall, both the SIT and MIT hypotheses predict faster reaction times (RTs) on target-match trials (i.e., only the target color matches one of the memory colors), and slower RTs on distractor-match trials (i.e., only the distractor color matches one of the memory colors). However, the SIT and MIT hypotheses make different predictions about what happens on individual trials. Specifically, when the target matches a VWM color, then the MIT hypothesis predicts that attention is always guided toward the target; in contrast, the SIT hypothesis predicts that attention is only guided toward the target on 50% of trials, because there is only a 50% chance that the target color serves as an attentional template.

Furthermore, we also manipulated the congruency between the target and the distractor to investigate whether both memory colors guide attention. Inside the target, the orientation of a line-segment was either congruent, or incongruent, with a line-segment inside the (colored) distractor. The MIT hypothesis predicts the strongest congruency effect on both-match trials (i.e., both the target and the distractor match the memory colors), because attention is simultaneously guided towards both the target and the distractor. Therefore, when the line-segment orientations of target and distractor are congruent, it is easier to report the orientation even though attention is partly drawn to the distractor, resulting in reduced RTs and error rates. In contrast, in the incongruent condition, there is more cognitive conflict caused by the different orientation of the matching distractor, resulting in increased RTs and error rates. The SIT hypothesis does not predict that attention is guided simultaneously towards the target and the distractor and therefore does not predict an especially strong congruency effect on both-match trials.

When it comes to the RT distribution of individual trials, the MIT hypothesis predicts that the distribution for both-match and non-match (i.e., neither the target nor the distractor match the memory colors) trials are the same, or at least very similar: On both-match trials, attention is guided toward both the target and the distractor, and the resulting facilitation and interference should approximately cancel each other out, resulting in an RT distribution that is similar to the condition where no color matches the VWM items. In contrast, under the SIT hypothesis, on both-match trials, attention is guided either toward the target, resulting in fast RTs, or toward the distractor, resulting in slow RTs, but never to both at the same time. Thus, the distribution for both-match trials is expected to be wider than that for non-match trials.^1^ We built drift-diffusion models of individual trials to simulate the two hypotheses’ predictions about RT distributions, and compared these with the collected data.

To foresee the results: The data by-and-large favor the predictions of the MIT hypothesis over the SIT hypothesis.

## Experiment 1

### Preregistration

Before conducting the experiment, we pre-registered the experimental designs on the Open Science Framework (OSF). A detailed pre-registration of the experiment is available at https://osf.io/sy7n8/. All deviations from the preregistration will be mentioned below.

### Method

#### Participants

We conducted a power analysis based on the results of a replication of Hollingworth and Beck (2016) as performed by Frătescu et al. (2019). Here the authors found that the effect size of the Distractor condition was *f* = 0.65. A power analysis conducted with G*Power (Faul et al., 2007) revealed that in order for this effect to be detected with a power of 95% and an alpha of .05, a sample of only seven participants would be required. Although this study is not identical to ours, this power analysis shows that memory-driven capture effects are strong and can be detected with few participants. However, our aim was to collect highly precise measurements that we could use also for computational modeling. In addition, we were interested in a modulation of the memory-driven capture effect by orientation congruency, and we had no a prior prediction about the strength of this modulatory effect. Therefore, we decided to collect at least 30 participants per experiment, which we felt confident would provide sufficient statistical power.

Thirty-five first-year psychology students (aged from 18 to 23 years old; 18 female, 17 male) from the University of Groningen participated in exchange for course credits. All participants had normal or corrected-to-normal acuity and color vision. The study was approved by the local ethics review board of the University of Groningen (18123-S). Participants provided written informed consent before the start of the experiment.

#### Stimuli, design and procedure

Participants were seated in a dimly lit, sound-attenuated testing booth, behind a computer screen on which the stimuli appeared at a viewing distance of approximately 62 cm. Stimuli were presented on a 27” flat-screen monitor at a refresh rate of 60 Hz running OpenSesame (version 3.2; Mathôt et al., 2012). Each trial started with a 500ms fixation display, followed by a 1,000ms memory display, consisting of two color disks (2.7° visual angle) placed in the middle of the screen to the left and the right of the fixation dot, with an eccentricity of 5.4° visual angle (*Figure 1*). The memory colors were randomly drawn from a HSV (hue-saturation-value) color circle with full value (i.e., brightness) and saturation for each hue (luminance ranged between 49 cd/m^2^ and 90 cd/m^2^), with the restriction that colors were at least 30° away from each other on the color circle. Participants were instructed to remember the exact colors of the items, and not the color category, to discourage verbalization.

**Figure 1.**
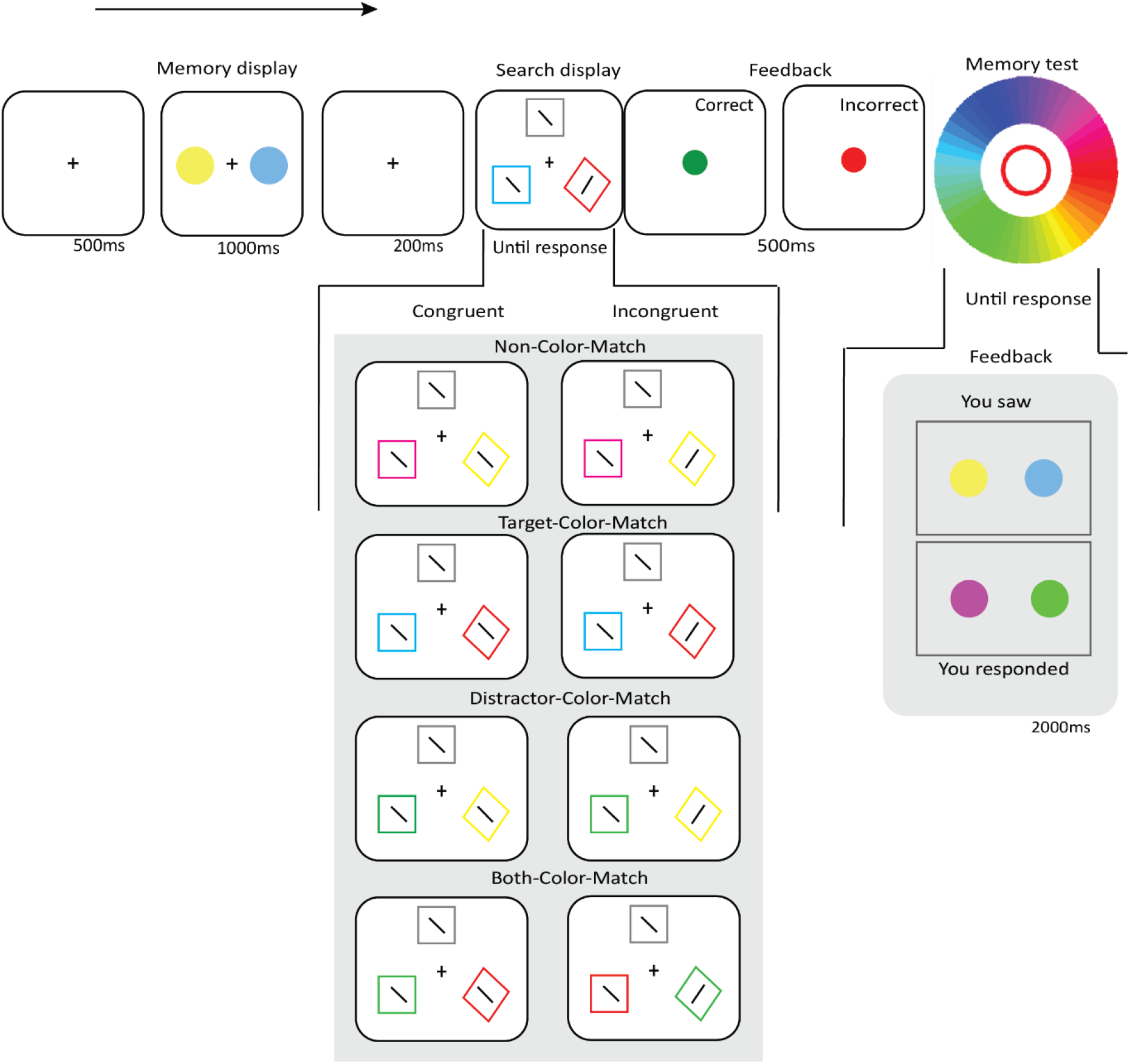
Sequence of events in a trial of *Experiment 1*. All Target-Color-Match and Distractor-Color-Match conditions in the search display for both Congruent and Incongruent trials are illustrated.

Following a 200ms fixation display, the search display was presented and remained visible until a response was given. The search display consisted of three shapes (1.3° visual angle): one diamond-shaped, colored target; one square-shaped, colored distractor and another square-shaped, gray distractor, all placed around the fixation dot, with an eccentricity of 5.4° visual angle). The colors of the target (diamond) and the colored distractor (square) either matched or did not match the remembered color depending on the *Target-Color-Match* (Match, Non-Match) and *Distractor-Color-Match* condition (Match, Non-Match), resulting in four types of trials: Non-Color-Match (i.e. target-color-non-match, distractor-color-non-match), Target-Color-Match (i.e., target-color-match, distractor-color-non-match), Distractor-Color-Match (i.e., target-color-non-match, distractor-color-match), and Both-Color-Match (i.e. target-color-match, distractor-color-match). All shapes in the search display contained a line segment (1.1° visual angle) that was tilted 22.5° clockwise or counterclockwise from a vertical orientation. The line segments in the target and the colored distractor were tilted in the same (Congruent) or a different (Incongruent) direction depending on the *Orientation-Congruency* condition. The line segment inside the grey distractor was chosen randomly, and will not be analyzed.

In our experiment, a color match was always exact; that is, when participants memorized a shade of green, then, on a Target-Color-Match trial, the visual-search target was always the exact same shade of green. However, this is not necessary for memory-driven capture to occur: both exact and inexact color matches lead to memory-driven capture (e.g., (Hollingworth & Beck, 2016); see also our own supplementary analysis on the OSF).

Participants indicated the orientation of the line segment within the diamond by clicking either the left or right mouse button as quickly and accurately as possible. Feedback was given for 500ms immediately following the response: a green dot for a correct response, or a red dot for an incorrect response. Each trial ended with a memory test, in which participants selected the exact color they memorized in the color circle. They did this twice, once for each memorized color. Visual feedback followed, comparing the colors they selected with those that they actually saw. The accuracy of each memory test was recorded as *memory precision*.

The three factors (*Target-Color-Match, Distractor-Color-Match, Orientation-Congruency*) were mixed randomly within blocks. Participants completed eight blocks of 32 trials each (256 trials in total), preceded by one practice block of 32 trials which was excluded from analysis.

#### Data processing

Trials with RTs shorter than 200ms and longer than 2,000ms were excluded. Next, participants were excluded from analyses if their accuracy on the search task was less than .7. (These criteria were not preregistered. We added them because our preregistered criteria failed to exclude some data points that were clearly unsatisfactory, such as participants who scored at chance level on the search task.) No participants were excluded based on our preregistered criterion of having a mean RT that deviated from more than 2.5 SD from the grand mean. Only RT data of correct trials were analyzed. Thirty participants and 7478 trials (of 8960) remained for further analysis.

#### Data analysis

The data were analyzed using the JASP software package (version 0.9; JASP Team, 2018) with the default settings, with *Target-Color-Match* (Match, Non-Match), *Distractor-Color-Match* (Match, Non-Match), and *Orientation-Congruency* (Congruent, Incongruent) as factors. (This deviates slightly from the preregistration, in which we treated Color-Match as a single factor with four levels.) We used inclusion Bayes Factor based on matched models (Rouder et al., 2009) to quantify evidence for effects.

Following (Lee & Wagenmakers, 2013), we considered Bayes factors (*BFs*) between 1 and 3 or between .3 and 1 as indicators of “anecdotal” evidence in favor of the alternative (*H_1_*) or the null hypothesis (*H_0_*), respectively; *BFs* between 3 and 10 or between .1 and .33 are indicators of “moderate” evidence; *BFs* between 10 and 30 or between .03 and .1 are indicators of “strong” evidence; and *BFs* between 30 and 100 or between .01 and .03 are indicators of “very strong” evidence of *H_1_* or *H_0_*.

### Results and Discussion

#### Search RTs

Analyses revealed very strong evidence for the effect of Target-Color-Match (BF_10_ = 3.30×10^24^) and Distractor-Color-Match (BF_10_ = 4.07 ×10^15^), such that RTs were faster when the target matched the memory color, and slower when the distractor matched the memory color (*Figure 2*). Moreover, we found moderate evidence for the effect of Orientation-Congruency (BF_10_ = 7.19), suggesting that RTs were faster on congruent trials than incongruent trials. No interaction effect between the factors was found (all BF_10_ < .06). (We also performed a supplementary analysis that included Memory Precision, based on a median split, as an additional factor. This revealed that memory precision of the VWM contents did not affect RTs or interact with any of the other factors. For more information, see the OSF project.)

**Figure 2.**
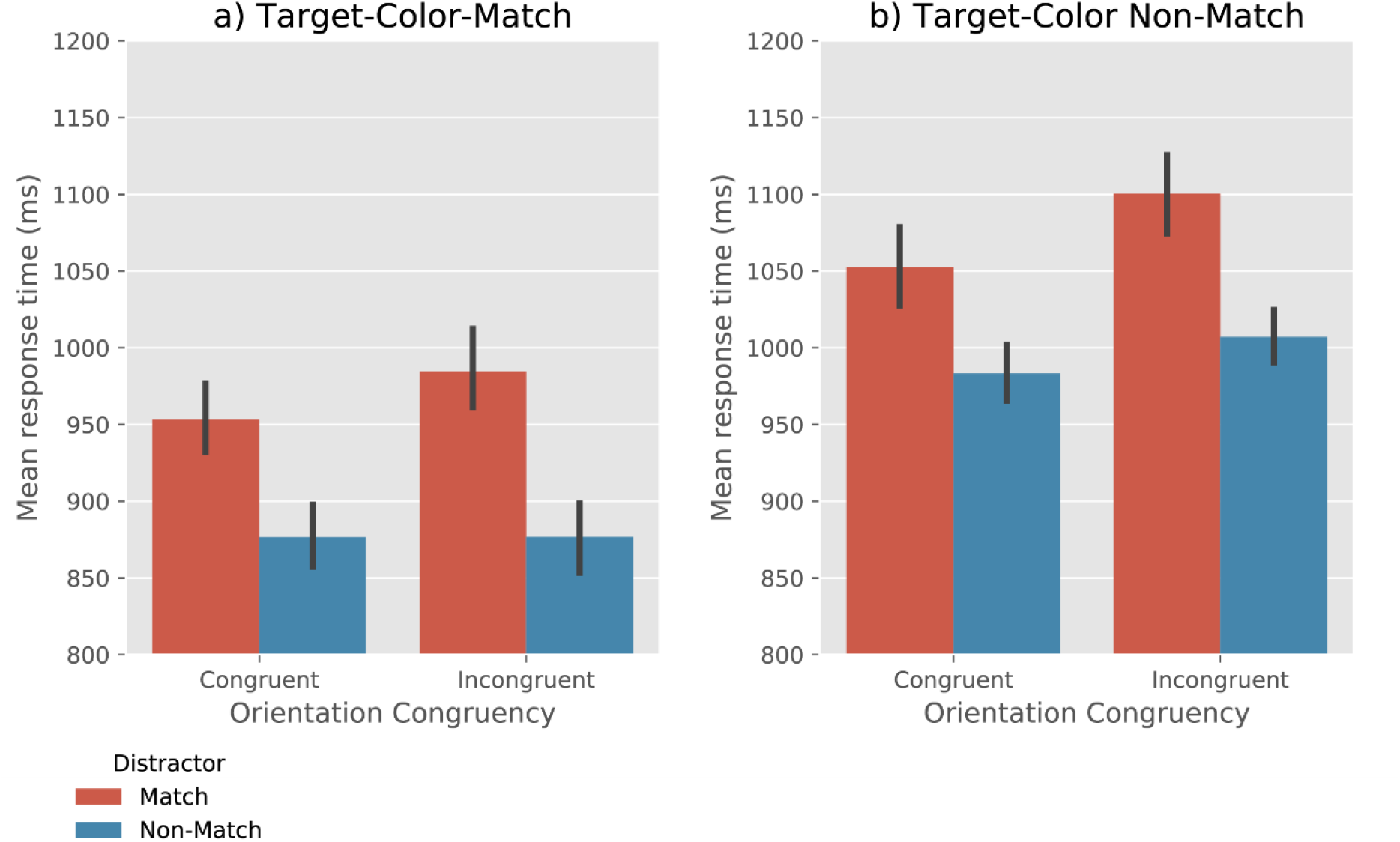
Mean response time as a function of Target-Color-Match, Distractor-Color-Match, and Orientation-Congruency. Error bars reflect condition-specific, within-subject 95% confidence intervals (Morey, 2008).

#### RT distributions

To test whether only one (i.e. SIT) or both (i.e. MIT) of the color items maintained in working memory served as attentional template, we analyzed the RT distributions for the Both-Color-Match and Non-Color-Match trials. According to the SIT hypothesis, on Both-Color-Match trials, attention is guided by the target on some trials, which leads to faster RTs, while on other trials attention is guided by the distractor, which leads to slower RTs. Therefore, the Both-Color-Match trials should result in a bimodal distribution (i.e. wider than that of the Non-Color-Match trials) according to the SIT hypothesis. In contrast, the MIT hypothesis predicts that on Both-Color-Match trials, both the target and the distractor guide attention, thus resulting in a unimodal distribution (i.e. resembling that of the Non-Color-Match trials).

To test this, an Inverse Gaussian distribution was fit to the RTs per condition for each participant. The scale parameter, which reflects the width of the distributions was analyzed using a evidenc T-test. We found moderate evidence that the RT distributions for the Both-Color-Match and the Non-Color-Match trials were equally wide (BF_01_ = 4.05, error % = .002), as predicted by the MIT hypothesis.

#### Accuracy

Analyses revealed moderate evidence for the effect of Target-Color-Match (BF_10_= 3.02) and Distractor-Color-Match (BF_10_ = 6.58), such that the overall search accuracy was higher when the target matches the memory color, and lower when the distractor matches the memory color (*Figure 3*). Furthermore, we found very strong evidence for the effect of Orientation-Congruency on accuracy (BF_10_ = 4.50×10^13^), showing that search performance was more accurate when the orientation of the line-segment in a target was congruent with that in a distractor than when they were incongruent. No evidence for any interaction effect between the factors was found (all BF_10_ < 2.0). (A supplementary analysis that included Memory Precision as an additional factor revealed that memory precision did not affect accuracy or interact with any of the other factors. For more information, see the OSF project.)

**Figure 3.**
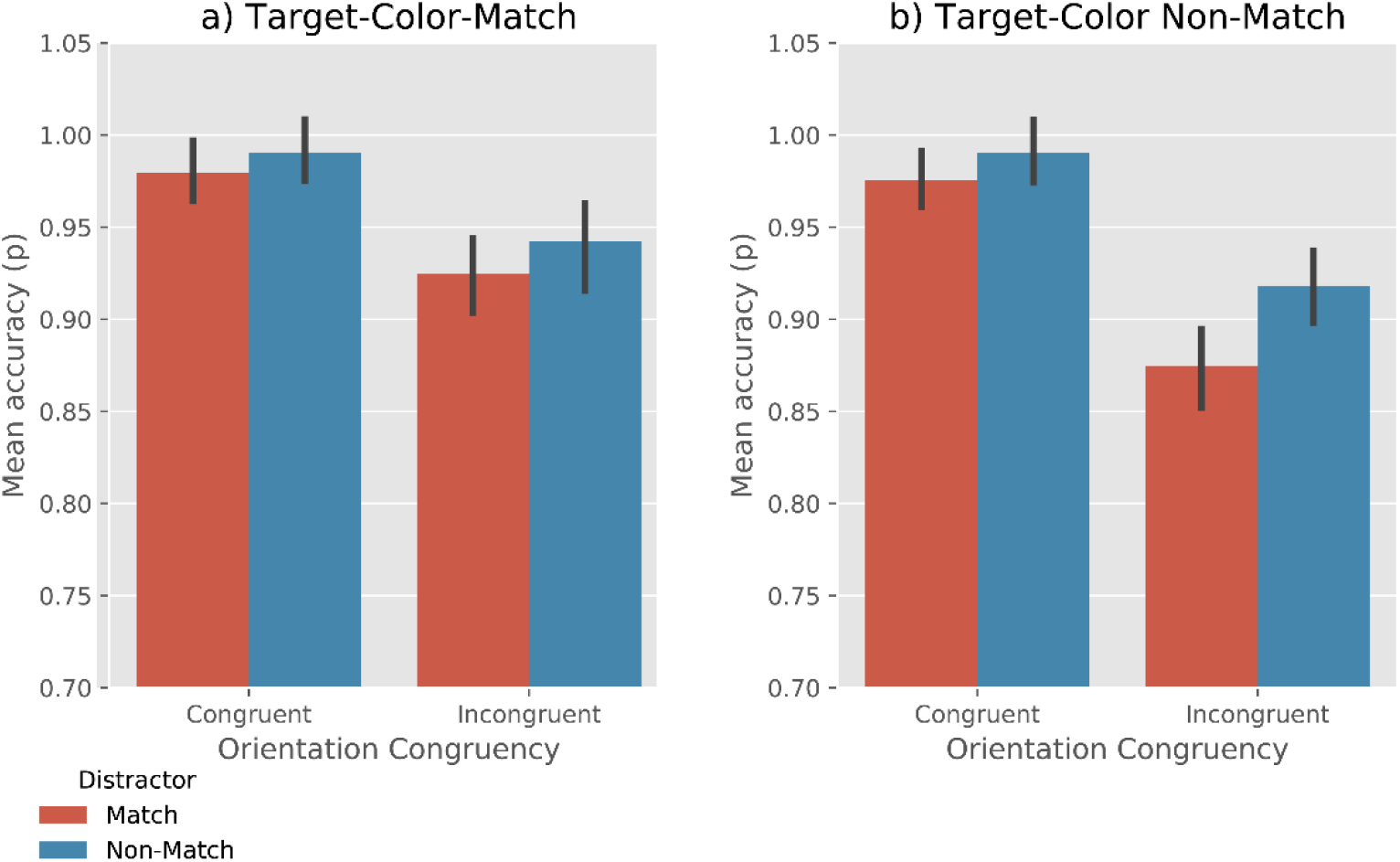
Mean accuracy rate as a function of Target-Color-Match, Distractor-Color-Match, and Orientation-Congruency. Error bars reflect condition-specific, within-subject 95% confidence intervals (Morey, 2008).

In summary, search performance increased (i.e. became faster and more accurate) when the target matched one of the colors held in VWM, but decreased when the distractor matched the VWM item. Moreover, the RT distribution for both-match trials and no-match trials are similar, which suggests that both color items that were maintained in the VWM draw attention. These results are consistent with the assumptions of the MIT hypothesis, as we will discuss in the General Discussion.

Unlike we predicted, however, we did not find that the effect of Orientation-Congruency was especially strong when both the target and the distractor matched, compared to other conditions. We suspected that the presence of the grey (unrelated) color might have affected the processing of the target and the distractor in visual search. Therefore, in the follow-up experiment, we removed the grey color in the search display.

## Experiment 2

In Experiment 2, we removed the grey color item (the unrelated item) from the search display. We reasoned that this would increase the strength of the Orientation-Congruency effect, because there were now only two line segments in the display, thus providing a stronger test of our prediction that the effect of Orientation-Congruency should be strongest when both the distractor and the target matched the VWM colors. Furthermore, we wanted to replicate the main results of Experiment 1.

### Preregistration

The preregistration was the same as in Experiment 1 expect for the data exclusion criteria, which now stated that the data would be trimmed based on a 70% accuracy rate. A detailed pre-registration of the experiment is available at https://osf.io/xpzhy/.

### Method

#### Participants

Thirty-six first-year psychology students (aged from 18 to 25 years old; 20 female, 16 male) from the University of Groningen participated in exchange for course credits. All participants had normal or corrected-to-normal acuity and color vision.

#### Stimuli, design and procedure

The method was the same as in Experiment 1 except for the following. The search display consists of one diamond-shaped, colored target, and one square-shaped, colored distractor, placed on an imaginary circle around the fixation with equal space between them (see *Figure 4*).

**Figure 4.**
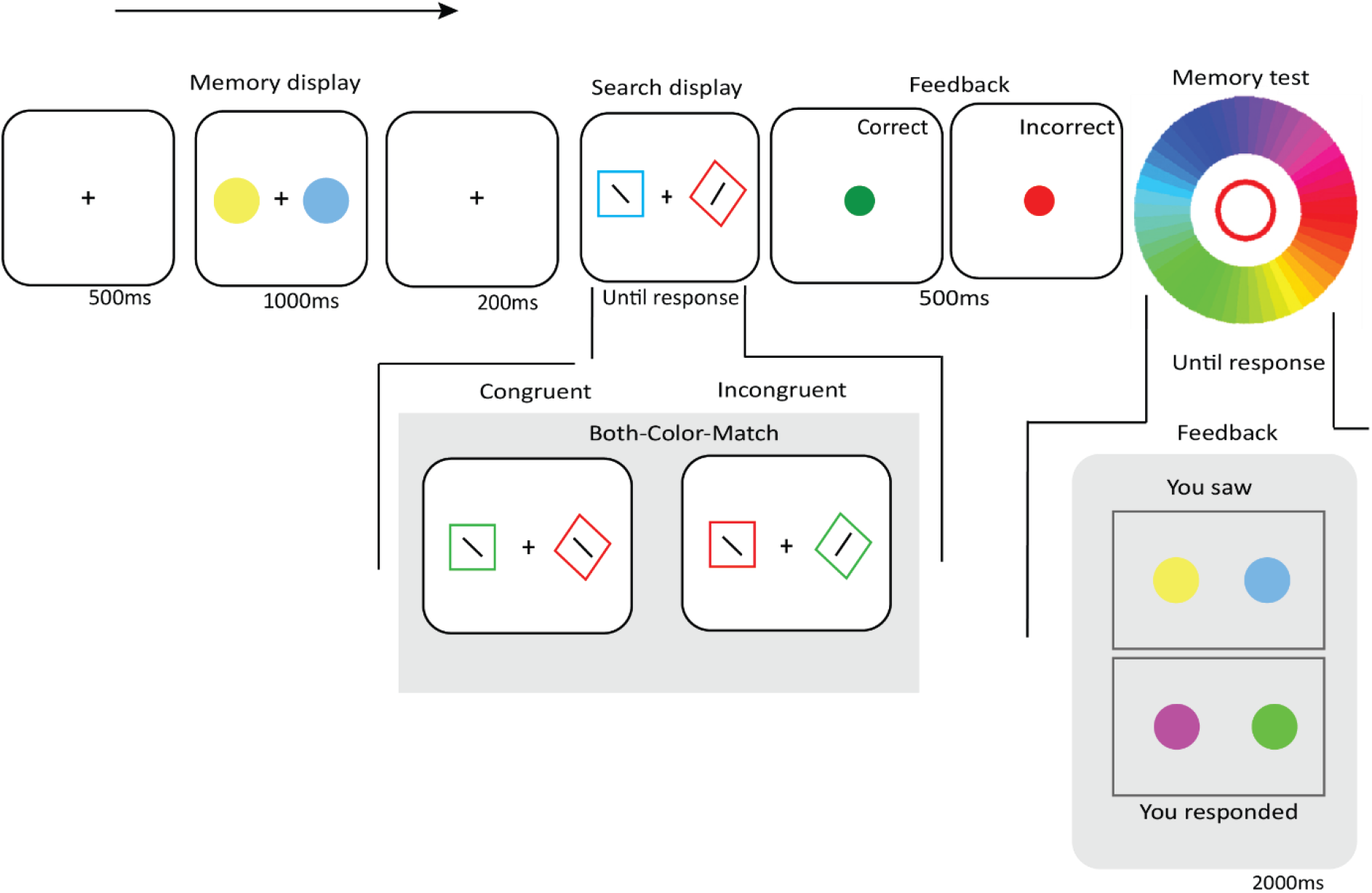
Sequence of events in a Distractor-Color-Match trial of *Experiment 2*.

#### Data processing

The same trimming criteria and analyses were used as in Experiment 1. Thirty participants and 7548 trials (of 9216) remained for further analysis.

### Results and Discussion

#### Search RTs

Analyses revealed very strong evidence for effects of Target-Color-Match (BF_10_= 2.15×10^6^) and Distractor-Color-Match (BF_10_ = 1.61×10^6^), such that RTs were faster when the target matched the memory color, and slower when the distractor matched the memory color (*Figure 5*). Moreover, we found a very strong effect of Orientation-Congruency on RTs (BF_10_ = 72.25), suggesting that participants were faster on congruent trials than on incongruent trials.

**Figure 5.**
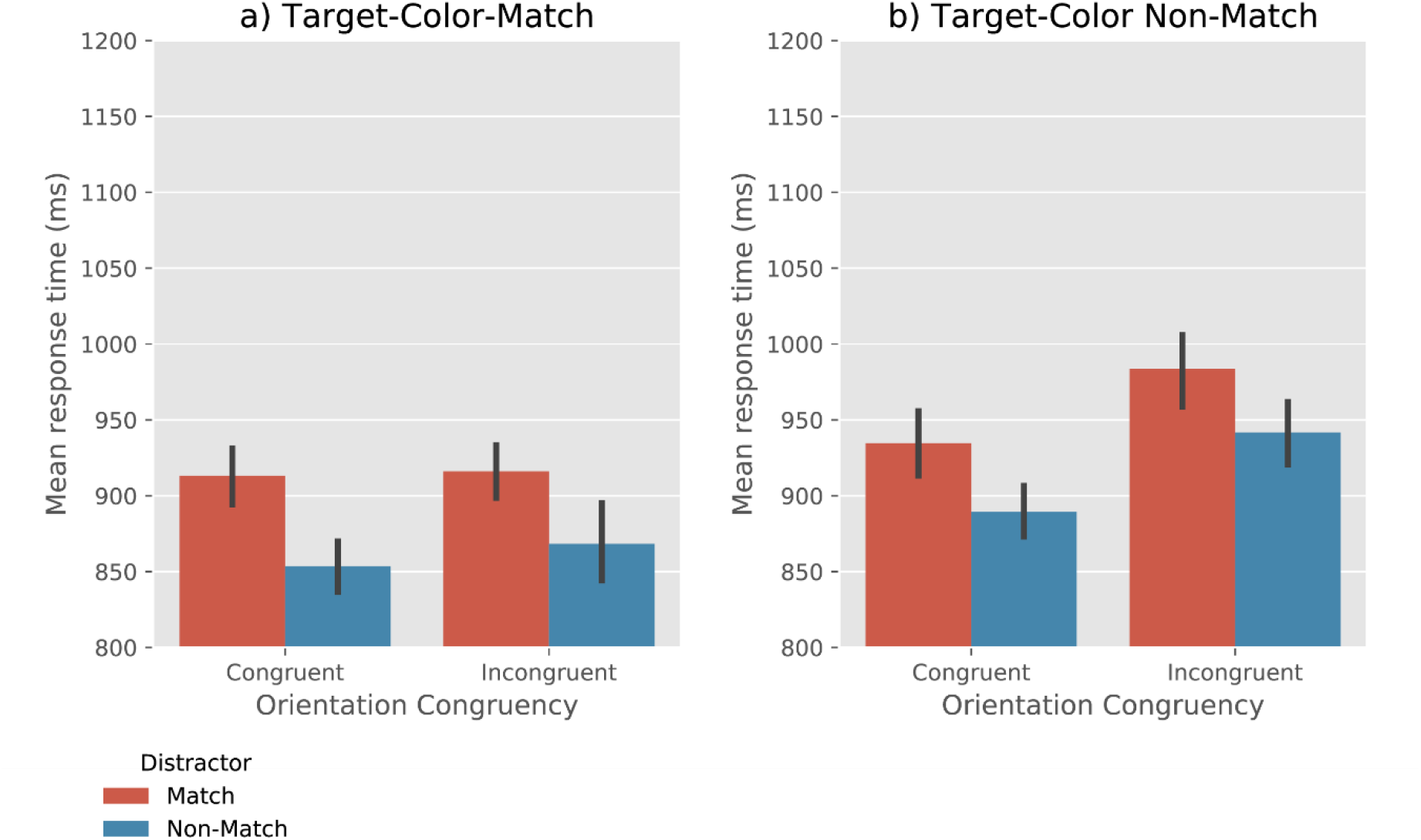
Mean response time as a function of Target-Color-Match, Distractor-Color-Match, and Orientation-Congruency. Error bars reflect condition-specific, within-subject 95% confidence intervals (Morey, 2008).

In addition, we observed moderate evidence for a Target-Color-Match × Orientation-Congruency interaction (BF_10_ = 3.69). To further qualify this effect, we performed a Bayesian ANOVA, with Orientation-Congruency and Distractor-Match as cofactors. When the target color did not match (*Figure 5b*), there was very strong evidence for a congruency effect (BF_10_ = 799.87); in contrast, when target matched the memory color (*Figure 5a*), there was moderate evidence *against* a congruency effect (BF_10_ = 0.245). No evidence for other interaction effects was found (all BF_10_ < .3). (A supplementary analysis revealed an effect of Memory Precision on RTs. This indicates that when the participants’ memory precision of the VWM items was higher, their search RTs were lower. There was no interaction of Memory Precision with any of the other factors. For more information, see the OSF project.)

#### RT distributions

Similar to Experiment 1, the RT distribution of the Both-Target-Match trials was equally wide as that of the Non-Match trials (BF01 = 5.14, error % = .01), as predicted by the MIT hypothesis.

#### Accuracy

Analyses revealed very strong evidence for effects of Target-Color-Match (BF_10_= 39.97) and Distractor-Color-Match (BF_10_ = 53.29), such that the accuracy was higher when the target matched the memory color, and lower when the distractor matched the memory color (*Figure 6*). Moreover, we found very strong evidence for the effect of Orientation-Congruency (BF_10_ = 1.19×10^6^), suggesting that search was more accurate when the line-segment orientation in a target was congruent with that in a distractor.

**Figure 6.**
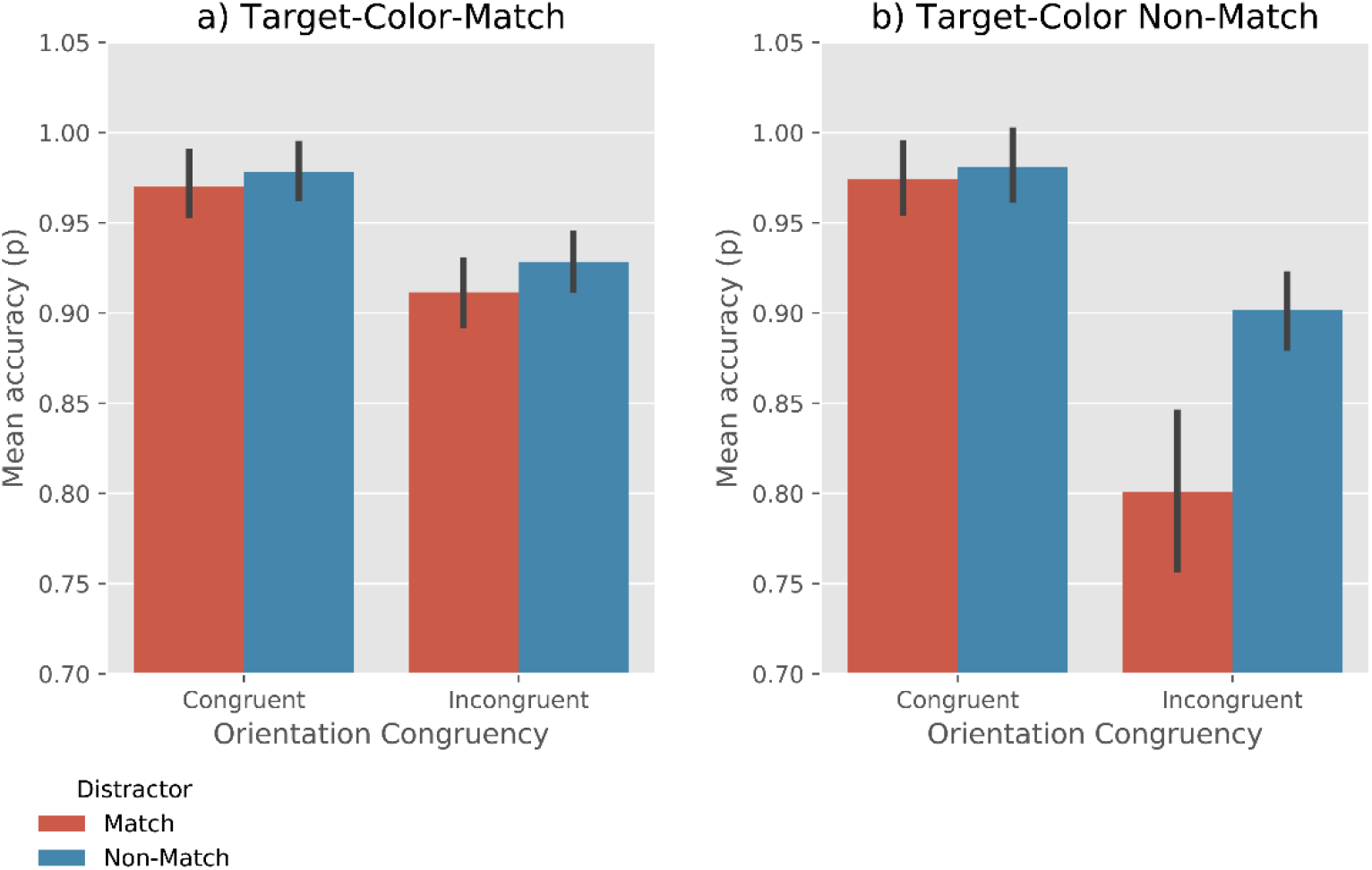
Mean accuracy as a function of Target-Color-Match, Distractor-Color-Match, and Orientation-Congruency. Error bars reflect condition-specific, within-subject 95% confidence intervals (Morey, 2008).

In addition, we observed a Target-Color-Match × Orientation-Congruency interaction (BF_10_ = 261.62). This indicates that the congruency effect was stronger when the target did not match the memory color. Furthermore, there was moderate evidence for Distractor-Color-Match × Orientation-Congruency interaction (BF_10_ = 8.15), suggesting that the congruency effect was stronger when the distractor matched the memory color. No reliable evidence for a three-way interaction (Target-Color-Match × Distractor-Color-Match × Orientation-Congruency) was found (BF_10_ = 2.74). (A supplementary analysis revealed an effect of Memory Precision on accuracy. This suggests that when the participants’ memory precision was high, their visual search more accurate. There was no interaction of Memory Precision with any of the other factors. For more information, see the OSF project.)

In this experiment, we observed faster overall RTs and stronger congruency effects than in Experiment 1. This suggests that the irrelevant (grey) distractor in Experiment 1 did attract attention, thereby reducing overall performance. Nevertheless, we successfully replicated the attentional guidance by the target and the distractor when they match the VWM colors. Moreover, we found that when the target matched the VWM item, the congruency effect largely disappeared; however, when the target did not match the VWM item but the distractor did match, the congruency effect was particularly strong. Although we did not predict this pattern of results, this robust guidance by the memory-matching item is in line with the MIT hypothesis, as we will discuss in the General Discussion.

## Drift-diffusion modeling

As described above, the distribution of correct RTs is very similar for the Non-Color-Match and Both-Color-Match trials; this favors the Multiple-Item-Template (MIT) hypothesis over the Single-Item-Template (SIT) hypothesis. However, we wanted to compare the predictions that both hypotheses make about RT distributions more rigorously.

To do so, we used a two-sided drift-diffusion model to simulate responses, and to generate error rates and distributions of correct RTs. The model simulates an Activation Level that changes over time, using four parameters: A Threshold, a Drift Rate, a Noise Level, and a Timeout. At time 0, the Activation Level is 0. At time 1, the Activation Level is incremented by the Drift Rate, as well as by a value that is randomly sampled from a normal distribution with a standard deviation that is equal to the Noise Level. Because we constrain the Drift Rate in our model to be a positive value, the Activation Level tends to increase over time, although with an element of randomness. The point in time at which the Activation Level reaches the threshold is taken as the simulated RT for a correct response; if the Activation Level reaches a value of minus the threshold, this is taken as an incorrect response. If the Activation Level has not reached a Threshold after a Timeout number of samples, the simulation is started again, until a valid RT is simulated. If no valid RT could be simulated after 1000 attempts, this was considered a failure to fit. A higher Drift Rate results, on average, in lower simulated RTs. A higher Noise Level results in more variable simulated RTs and increased error rates.

The Threshold was set to a constant value of 1. The Timeout was set to a constant value of 2000, corresponding to the 2000ms timeout in our experiments. The Drift Rate and Noise Level were determined for each participant separately, by taking all the RTs for a given participant, and rank-ordering them first based on whether they were correct or not, and then based on their value. Next, we simulated the same number of correct and incorrect RTs, using a candidate pair of values for the Drift Rate and Noise Level, and similarly rank-ordered these simulated RTs. We then took the residual sum of squares (RSS) of the real and simulated RTs. The Drift Rate and Noise Level were then chosen such that they minimized the RSS for a given participant. Phrased differently, we chose parameters such that they minimized the error between the real and simulated RT distributions for both correct and incorrect responses.

Next, we constructed two models that embodied the predictions of the MIT and SIT hypotheses. To do so, we added one additional parameter, Drift Rate Change, which was added to the basic Drift Rate to simulate the reduced RTs (facilitation) when attention was guided by the Target, and subtracted from the basic Drift Rate to simulate the increased RTs (interference) when attention was guided by the Colored Distractor. To keep the number of model parameters to a minimum, we used a single parameter for the Drift Rate Change for both facilitation and interference, rather than two separate parameters. This choice reflects our assumption that facilitation and interference should approximately cancel each other out, although there is no theoretical reason to assume that they do so perfectly.

The MIT and SIT hypotheses make slightly different predictions about the Drift Rate in the different conditions (Table 1). In a nutshell, the MIT hypothesis predicts that a Target-Color-Match should result in facilitation on every trial, and that a Distractor-Color-Match should result in interference on every trial, and that the two should approximately cancel each other out on both-match trials. In contrast, the SIT hypothesis predicts that a Target-Color-Match should result in facilitation on only 50% of trials, because only one of the two VWM items serves as an attentional template, and thus the probability of the Target matching the attentional template is only 50%. For the same reason, a Distractor-Color-Match should result in interference on only 50% of trials, and the Both-Color-Match condition should be a mixture of 50% facilitation and 50% interference.

**Table 1.** The Drift Rate in each condition as predicted by the MIT and SIT models. The percentages indicate the percentage of trials on which the Drift Rate has a particular value.

| **Trial Type** | **MIT model** | **SIT model** |
| --- | --- | --- |
| Non-Color-Match | 100%: Drift Rate | 100%: Drift Rate |
| Target-Color-Match | 100%: Drift Rate + Drift Rate Change | 50%: Drift Rate + Drift Rate Change<br>50%: Drift Rate |
| Distractor-Color-Match | 100%: Drift Rate - Drift Rate Change | 50%: Drift Rate - Drift Rate Change<br>50%: Drift Rate |
| Both-Color-Match | 100%: Drift Rate | 50%: Drift Rate + Drift Rate Change<br>50%: Drift Rate – Drift Rate Change |

For each participant separately, and for the MIT and SIT models separately, we then determined the Drift Rate Change parameter, while keeping the other parameters as previously determined. This was done by taking all the RTs for a given participant, ordering them first by whether they were correct or not, then by trial type (Non-Color Match, Target-Color-Match, Distractor-Color-Match, Both-Color-Match), and then rank-ordering them from fast to slow. We then simulated the same number of RTs, using a candidate value for the Drift Rate Change, and similarly ordered these simulated RTs. The Drift Rate Change was then chosen such that it minimized the RSS between the real and simulated RTs. For the SIT model (but not the MIT model), even the optimal parameters failed to generate a sufficient number of incorrect responses for twelve participants; these participants were excluded from the analysis below, although these failures-to-fit already illustrate that the SIT model is less able to characterize human data than the MIT model is.

To test which model could best account for the data, we compared the RSS for the MIT model and the RSS for the SIT model with a default Bayesian, as well as a traditional, two-sided paired-samples t-test. This revealed very strong evidence (BF_10_ = 524; error % = 2.67×10-10; t(47) = 4.52, p < .001) in favor of the MIT hypothesis. To qualitatively compare the MIT and SIT model to the human data, we generated distributions of correct RTs, which were z-scored for each participant for visualization, as well as error rates. As shown in Figure 7, the MIT model characterizes the human data better than the SIT model does, both in terms of correct RTs and error rates.

**Figure 7.**
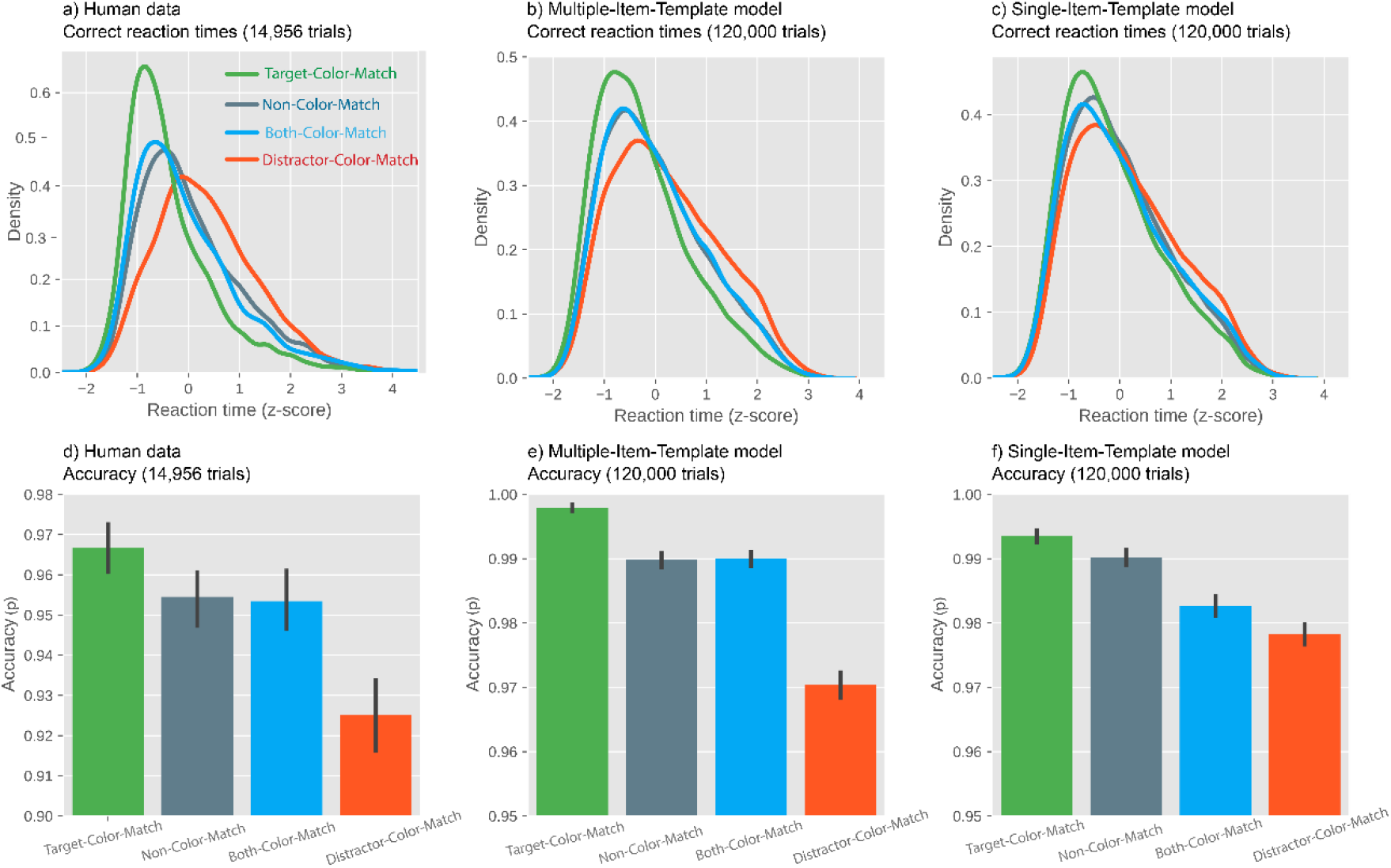
Top row: Distributions of correct response times for a) human data, b) the Multiple-Item-Template (MIT) model, and c) the Single-Item-Template (SIT) model. Bottom row: Accuracy (proportion of correct responses) for d) human data, e) the MIT model, and f) the SIT model.

## General Discussion

Here we report that multiple working-memory representations guide attention concurrently, thus providing crucial behavioral and computational evidence for a long-standing debate in the field of visual working memory (VWM). In our experiments, participants remembered two colors. Next, they performed a visual-search task in which the color of the target and that of a distractor could match, or not match, a color in VWM. We found that search was faster when there was a target-color match, showing that attention was guided towards memory-matching targets; similarly, we found that search was slower when there was a distractor-color match, showing that attention was (mis)guided towards memory-matching distractors.

To further test the predictions of the Multiple-Item-Template (MIT) and Single-Item-Template (SIT) hypotheses, the orientation of the line-segment inside the search target was manipulated to be either the same (i.e., congruent) or opposite (i.e., incongruent) to the line segment inside the distractor. Overall, this should result in an Orientation-Congruency effect, such that RTs are slower on incongruent compared to congruent trials if attention is divided between the target and the distractor. However, the MIT and SIT hypotheses make different predictions about when this congruency effect should be strongest. Specifically, the MIT hypothesis predicts that the congruency effect should be strongest on both-match trials (i.e. when both the target and the distractor matched the memorized colors). This prediction follows because only in that case attention would be drawn simultaneously towards the distractor and the target, thus creating the strongest interference (and thus the strongest congruency effect) in that condition. The SIT hypothesis makes no such prediction, because on both-match trials, attention would be guided either by the target or by the distractor dependent on which of the colors was used as a template color, but not by both, and thus there is no reason to predict increased interference.

Although, we did *not* find an increased congruency effect on both-match trials, we *did* find that the congruency effect was largely absent whenever there was a target match in Experiment 2. This implies a two-stage model of visual search (Kastner & Nobre, 2014). First, attention is guided in parallel to (the color of) all memory-matching stimuli, resulting in facilitation by matching targets, and interference by matching distractors; that is, activation in the priority map is affected by the content of VWM. Next, the orientations of the line-segments inside the stimuli are processed serially; that is, highly activated items in the priority map are further processed one after another. On target-match trials, the line-segment inside the target is generally processed first, because participants have a search template for the target’s shape (a diamond), which gives the target additional activation in the priority map; next, once the target has been processed, a decision is made, and the line-segment inside the distractor is left largely unprocessed. This would explain the strongly reduced interference by incongruent distractors on target-match trials. In general, this finding suggests that, on most trials, attention was captured by the memory-matching target (and not only on 50% of trials). Although we did not predict this, it is consistent with the MIT hypothesis that two templates can be simultaneously activated to guide attention. Compared to Experiment 1, in Experiment 2 we removed the unrelated distractor (i.e. the grey colored item) to reduce attentional capture by non-relevant distractor items, thereby inducing a stronger congruency effect, thus changing the task from a regular visual-search task to a discrimination task between a target and a single (colored) distractor. Crucially, the results remained qualitatively the same, suggesting that our results do not depend on the specifics of the task. Nevertheless, future studies could explore how including more search elements (e.g., more colored distractors that never match) affects the pattern of results.

In our paradigm, whenever there was a match between a memorized color and the color of an item in the search task, this match was always perfect. This raises the possibility that participants strategically attended to matching targets and distractors, to refresh their memory. However, previous studies have shown that memory-driven guidance of attention also occurs when there is only a categorical match (e.g. when participants memorize a shade of green, and the search distractor is a slightly different shade of green; Hollingworth & Beck, 2016; replicated in Frătescu et al., 2019). Therefore, our results are unlikely to depend on the use of perfect color matches. Nevertheless, the flexibility of memory-driven guidance is an important direction for future research: what exactly does it mean for visual input to ‘match’ the content of VWM?

Additionally, we analyzed the RT distribution for both-match and no-match trials (i.e. when neither the target nor the distractor matched the memorized colors). The MIT hypothesis predicts that the distribution for both-match and no-match trials should be the same (or at least similar). This follows from the MIT hypothesis, because on both-match trials, the facilitation due to attention being guided towards the target and the interference due to attention being guided towards the distractor should approximately cancel each other out. In contrast, the SIT hypothesis predicts a wider distribution for both-match trials than for no-match trials. This follows from the SIT hypothesis, because attention is guided either by the target or by the distractor in both-match trials (but never by both), thus resulting in a bimodal distribution that is wider than the distribution for no-match trials. Consistent with the MIT hypothesis, we found that the RT distribution for both-match trials resembled that for no-match trials. This implies that not only can multiple VWM items serve as attentional templates, but that it is also possible for focal attention to be allocated to multiple items at the same time rapidly (Eimer & Grubert, 2014). To confirm this conclusion, we simulated the individual trials of RTs based on the predictions of the MIT and the SIT hypothesis by means of a drift-diffusion model. Crucially, the observed data showed a better match to the simulated RTs based on the MIT hypothesis.

Taken together, our results provide evidence against the SIT hypothesis, which posits that there can only be one template active in working memory at one time to bias visual selection (Olivers et al., 2011; van Moorselaar et al., 2014). And we show behavioral and computational evidence for simultaneous guidance of multiple VWM items, providing support for the MIT hypothesis.

## Open Practices Statement

All experimental data and materials can be found on the OSF (Open Science Framework): https://osf.io/knmu2/. The pre-registrations of the experiments are available at https://osf.io/knmu2/registrations.

## Footnotes

1 One can also imagine a version of the MIT hypothesis in which guidance of attention is parallel (such that multiple items can draw attention), but that due to a winner-takes-all process, attention is only deployed to a single item at a time (i.e. deployment of attention is serial). This model, which we will not consider further, makes predictions that are very similar to the SIT model, and for the present discussion can be considered analogous to the SIT hypothesis.

